# SNVstory: A dockerized algorithm for rapid and accurate inference of sub-continental ancestry

**DOI:** 10.1101/2023.06.02.543369

**Authors:** Audrey E. Bollas, Andrei Rajkovic, Defne Ceyhan, Jeffrey B. Gaither, Elaine R. Mardis, Peter White

**Author notes:** Corresponding Author: Peter White, Ph.D., The Institute for Genomic Medicine The Abigail Wexner Research Institute Nationwide Children’s Hospital, 575 Children’s Crossroad, Columbus, OH 43215.

## Abstract

Knowing a patient’s genetic ancestry is crucial in clinical settings, providing benefits such as tailored genetic testing, targeted health screening based on ancestral disease-predisposition rates, and personalized medication dosages. However, self-reported ancestry can be subjective, making it difficult to apply consistently. Moreover, existing approaches utilize genome sequencing data to infer ancestry at the continental level, creating the need for methods optimized for individual ancestry assignment. We present SNVstory, a method built upon three independent machine learning models for accurately inferring the sub-continental ancestry of individuals. SNVstory includes a feature-importance scheme, unique among open-source ancestral tools, which allows the user to track the ancestral signal broadcast by a given gene or locus. We apply SNVstory to a clinical dataset, comparing self-reported ethnicity and race to our inferred genetic ancestry. SNVstory represents a significant advance in methods to assign genetic ancestry, predicting ancestry across 36 different populations with high accuracy.

## Introduction

Ancestry derived from genomic data, referred to as genetic ancestry, is a measurable and biologically defined parameter. Although much of the human genome is identical across all populations, it is estimated that depending on an individual’s ancestry, 0.1% to 0.4% may differ from the human reference genome. While this genetic variation includes structural variants (SVs), copy number variants (CNVs), and small insertions or deletions (indels), by far the largest and easiest to detect category occurs in the form of single nucleotide variants (SNVs), many of which are unique to genetically distinct populations^1^.

Knowledge of a patient’s genetic ancestry has clinical implications, ranging from genetic testing to health screening based on ancestral disease-predisposition rates, and in some cases, may inform what medicine dosage to prescribe a patient^2–4^. However, self-reported race is frequently used in the research and clinical setting and is often inconsistent with genetic ancestry, potentially driving health disparities^5–8^. Genome sequencing-based diagnostic testing in patients suspected of having a rare genetic disorder requires accurate data filtering to remove variants common to a given population. Precise identification of the patient’s ancestry improves the identification of rare disease-causal variants. Therefore, developing methods to report ancestry accurately and consistently is essential.

In addition to clinical importance, knowing the ancestral composition of an individual or a population is essential in the genetic research setting. For example, signals from genome-wide association studies (GWAS) or whole genome sequencing cohorts can be reassessed based on population stratification, whereby loci associated with disease may be more accurately identified by discarding rare variants associated with an individual’s ancestry rather than with the disease in question^9,10^.

Given the importance of ancestry, several ancestry inference algorithms that operate on genomic data have been developed that can be divided into two broad types: parametric and non-parametric. Parametric learning algorithms estimate a finite set of parameters from the data to establish a relationship between the independent and dependent variables. Two widely-used parametric tools are STRUCTURE^1^ and ADMIXTURE^12^, which estimate the proportions of different ancestries (or ancestral populations) for each individual, known as admixture. Recently, Archetypal Analysis was shown to be more computationally efficient and provide more interpretable results than ADMIXTURE^13^. In contrast, non-parametric methods do not have a finite set of parameters and instead rely on the intrinsic structure of the data to determine which data points best resemble each other.

The emergence of population-scale genome sequencing datasets with a form of self-reported ancestry allows models to be built with prior knowledge of represented ancestries. In place of individualized genetic data, large databases house genomic summary results, such as aggregate variant allele frequencies stratified by population. For example, the Single Nucleotide Polymorphism database (dbSNP) is the largest genomic aggregate database with 11 different populations from over one million samples^14^. However, the 11 distinct populations contain a high degree of overlap and primarily represent continental groupings^15^. The Genome Aggregation Database (gnomAD) is another aggregate database with allele frequencies from 140,000 subjects from 26 populations^16^. In addition to these large-scale repositories of aggregate allele frequencies, there exist a few datasets at the level of the individual, such as the 1000 Genomes Project (1kGP)^1^ and the Simons Genome Diversity Project (SGDP)^17^, which are much smaller in sample size, with 2,504 and 279 samples, respectively. Nevertheless, the 1kGP and SGDP have been critical in characterizing ancestry and human history as they contain the most granular population labels.

Taken together, these curated variant datasets enable an alternative class of models to be used to predict ancestry based upon samples labeled with known ancestry^18–28^. However, many methods suffer shortcomings, including not having discrete ancestry labels beyond the main continental groups or, for those methods using the 1kGP, not considering that many subjects are within the same families and, therefore, fail to satisfy the principle of independent and identically distributed data. As such, there is a critical need for methods to accurately predict an individual’s genetic ancestry from genome sequencing data by implementing supervised models.

Here, we address some limitations surrounding supervised learning of ancestry by developing three independent models from gnomAD, 1kGP, and SGDP. Our models estimate ancestry from 36 different populations with high accuracy. Furthermore, we provide software that enables users to run our models on their data, taking the widely accepted variant call format (VCF) files as input and outputting predictions and a graphical representation of the likelihood of a given genetic ancestry. As a form of validation, we apply these models to our in-house clinical research dataset and correlate the estimates with those of self-reported ancestry.

## Materials and Methods

### Training Datasets

Genomic datasets from gnomAD, 1kGP, and SGDP were processed separately (**Figure 1**), as described below. The gnomAD variants are provided on reference genome GRCh37, and the 1kGP and SGDP were called on reference genome GRCh38.

**Figure 1.**
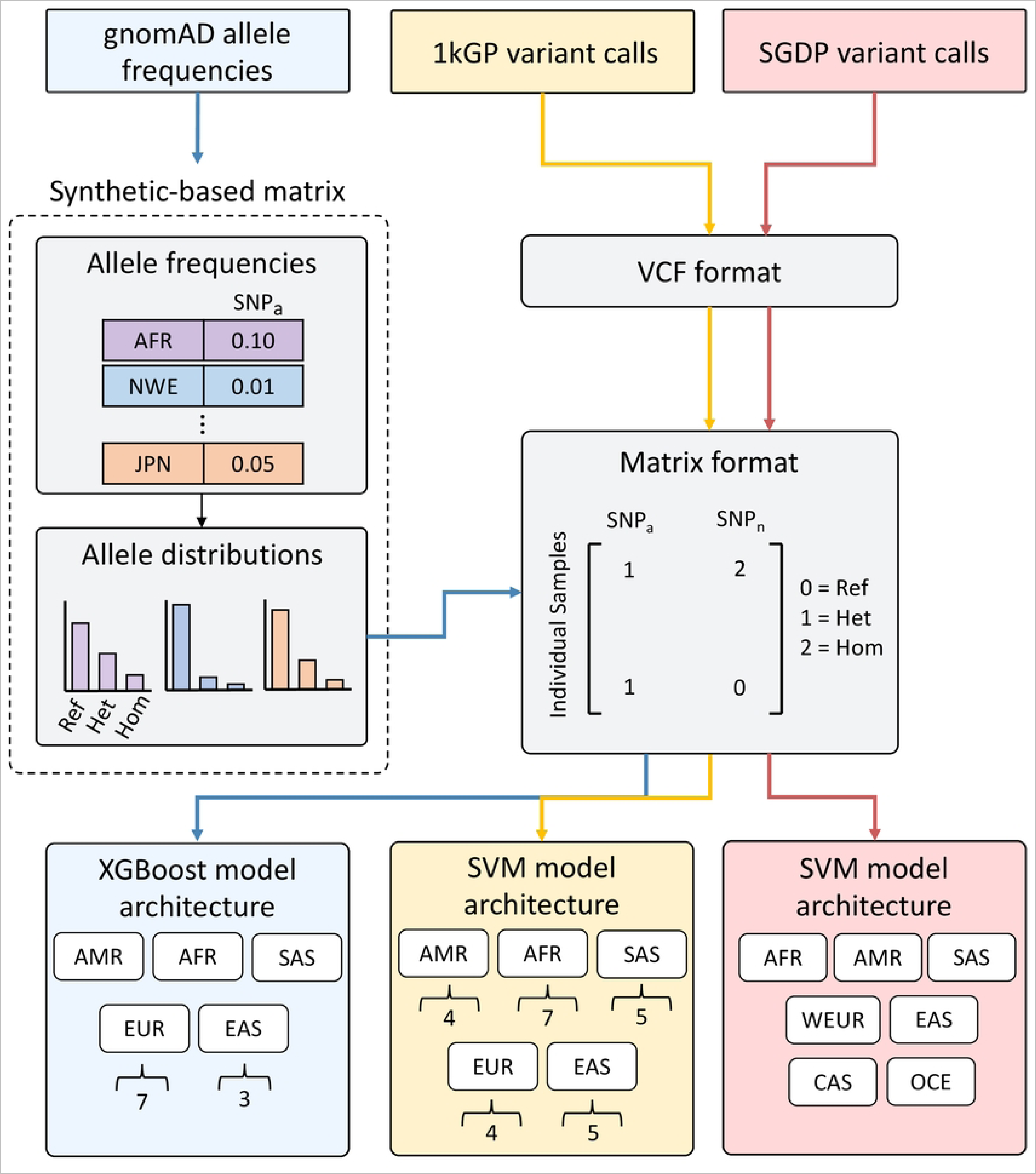
Schematic of ancestry inference model strategy. The workflow visualizes each dataset separately with colored boxes and arrows: gnomAD (blue), 1kGP (yellow), and SGDP (red). For the gnomAD synthetic-based matrix, allele frequencies for each variant for each population given in gnomAD are used to create a distribution of reference, heterozygous and homozygous alleles for each population. A matrix format is created by converting the distributions into 0’s, 1’s, and 2’s for each locus for samples in each population. For 1kGP and SGDP, a matrix format is built directly from variants in the VCF. For the model architecture, continental model labels are shown in white boxes, and the number of labels in the corresponding subcontinental models is below in brackets.

### The Genome Aggregation Database (gnomAD)

The gnomAD v2.1 exome and genome sequencing variant dataset provides aggregated data from 17 populations, meaning allele frequencies of each population for 17 million exome variants. We reduced the number of input features for machine learning by following a similar protocol to the one described by the MacArthur lab by filtering for high call rates, biallelic-only sites, and a frequency greater than 0.1% (https://macarthurlab.org/2018/10/17/gnomad-v2-1/). After this filtering, 81,398 SNVs remained, formatted as a matrix of ancestries and corresponding SNV frequencies.

To obtain SNV calls for individuals, as is provided in standard VCF format, we simulated individuals from each ancestry by effectively flipping a weighted coin for each individual and their respective variant (**Figure 1**). This resulted in a synthetic-based matrix of samples spanning the ancestry classifications in gnomAD v2.1 and SNVs, coded as reference, heterozygous, or homozygous for each SNV position. Although this approach does not capture haplotypes, the simulated samples are genetically typical examples of the chosen ancestry to a first approximation.

### The 1000 Genomes Project (1kGP)

The New York Genome Center performed genome sequencing (GS) on 3,202 samples, including 602 trios, from the 1kGP cohort at 30x coverage, released in 2020^29^. The data were aligned to GRCh38 using BWA-MEM^30^, and variants were called by GATK *HaplotypeCaller* (GATK version 3.5.0) using default settings. The dataset contains 126,659,422 SNVs from 26 populations spanning East and South Asia, North and South America, Africa, and Europe. Sample sizes were not uniformly represented across the different populations, i.e., the dataset was imbalanced. Due to the high genetic similarity between individuals from Utah and the United Kingdom, the Utah population was removed from the analysis.

### The Simons Genome Diversity Project (SGDP)

The SGDP consists of GS of 300 individuals from seven major population groups, 75 countries, and 142 diverse populations. GS FASTQ files from 279 samples were downloaded from the European Nucleotide Archive (PRJEB9586). Sequencing reads were aligned to genome assembly GRCh38 using BWA-MEM. SNV and INDEL calling was performed with GATK version 4.1.9, described below. GATK *HaplotypeCaller* was run on each sample using the GVCF workflow to generate a per-sample intermediate GVCF. The GATK *GenotypeGVCFs* function was used to perform base calling across all samples jointly to obtain genotypes for each sample in VCF format. We then performed variant recalibration and filtering in the two-stage process using the GATK functions *VariantRecalibration* and *ApplyVQSR*. The final combined data set contained a total of 48,815,712 SNVs.

### Quality Control

Quality control of the gnomAD (https://macarthurlab.org/2018/10/17/gnomad-v2-1/) and 1kGP^29^ were as previously described. For the SGDP dataset, we ran several quality-control tools to detect any issues with sequencing quality and sample contamination. We ran Picard *CollectMultipleMetrics* on the aligned bam files to collect alignment summary, quality score, and GC bias metrics (**Table S1**). Sequencing read allocation was calculated using samtools. Coverage information was collected using mosdepth^31^. The average coverage for all realigned samples was 40X (ranging from 31X to 77X). Sample contamination level was determined by the number of reads inconsistent with the genotype in dbSNP^14^ sites. One sample was flagged for possible sample contamination (**Supplemental Materials and Methods**).

### Removal of Related Samples

Related samples of the third degree (e.g., first cousins, great grandparents, or great-grandchildren) or closer were identified by the relationship inference tool, KING^32^. Data from the 1kGP and SDGP were preprocessed using PLINK2 with the following parameters: *“--new-id-max-allele-len 10000 -- max-alleles 2*”^33^. KING recommends performing as little filtering as possible. However, an additional filtering step was performed to prevent the computation from running out of memory. Therefore, the analysis was restricted to variants shared by at least two individuals: “*--maf 0.0007*” in the case of the 1kGP and “*--maf 0.007*” for SDGP. After removing the variants present in only one sample, KING was executed on the resulting bed file, with the “*--kinship*” option set to report pairwise relatedness inference. Samples from the analysis were flagged that had a third-degree kinship coefficient cutoff >= 0.0442, a value previously established by the authors of KING^32^. Four samples were removed from further analysis in the SGDP dataset based on the KING relatedness results (**Supplemental Materials and Methods**).

Because some samples from the 1kGP are related to more than one other individual in the cohort, the following procedure was implemented to remove the fewest number of samples. Considering only the relationships with coefficients exceeding the third-degree cutoff, a graph-based method was implemented to recursively identify nodes (samples) with the largest number of edges (relationships) and remove those nodes until all subgraphs had, at most, a single connection. For subgraphs with a single connection, one sample was randomly selected from the pair, while all singletons were included in the list of samples to keep. From 167 samples with at least one close relationship, 117 were flagged for inclusion in downstream analysis. The remaining samples were removed with PLINK2.

### Variant Selection and Preprocessing

Variants from 1kGP and SGDP underwent a final filtering step by taking the intersection of targeted exonic regions of the exome capture reagent used routinely in our clinical lab (IDT xGen Exome Hyb Panel v2 targets hg38 BED file) with the set of genetic variants from the unrelated individuals using BEDTools *intersect* (v2.30.0)^34^. The resulting VCF was converted into a numerical encoding homozygous alternative = 2, heterozygous = 1, reference or missing = 0. The vectors of genotypes were combined to form a matrix of variants by genotypes. For variant selection from gnomAD, see the following gnomAD section in Model training and cross-validation below.

### Model Training and Cross-Validation

The models were trained on each dataset separately, as required by their differing labeling strategies (**Figure 1**).

#### gnomAD

Because our gnomAD algorithm uses synthetic data, we must consider two parameters: a population size that balances the model’s accuracy with training time and resources and a p-value from a Chi-Square test that removes uninformative SNVs. This was accomplished using a nested for loop to iterate over all combinations of population sizes and p-values for SNV removal (**Figure S1**). For each combination, we generated a set of 80/20 training/validation splits of the data. A Chi-Square test was applied to each SNV (feature) in the training data to determine whether it was informative for distinguishing ancestry in the population. SNVs were removed that did not meet the p-value threshold. We used a gradient-boosted decision tree from XGBoost to train the model on the training set and then test on the validation set^35^. Fold generation and training were performed five times for each p-value, and the accuracy was averaged to represent the accuracy for each p-value. Once all the p-values were tested, the p-value with the highest accuracy was selected (**Figure S2**). Then, the model was retrained on all the data for that specific population size and tested on a synthetic hold-out set. The accuracy for the hold-out set is representative of that population. A continental model (population size of 4,084 individuals; SNV p-value threshold of 7.5e-49) was built to predict six groups: Africa, South Asia, Europe, East Asia, America, and Ashkenazi Jewish. Two sub-continental classifiers were built to predict ancestry within the East Asian (**Figure S2A.**; population size of 13,593 individuals; SNV p-value threshold of 1.78e-09) and European groups (**Figure S2B.**; population size of 45,243 individuals; SNV p-value threshold of 1.78e-24).

#### 1kGP

For the 1kGP dataset, the support vector machine (SVM) library from scikit-learn^36^ was used to train a classifier to predict the continental groups: Africa, Europe, South Asia, East Asia, and America. In addition, multiple classifiers were trained independently for each sub-continental group, i.e., Kenya or African Caribbean in Barbados. All SVMs were trained using the radial basis function (RBF) kernel and with the gamma parameter fixed as the default. Hyperparameter tuning of the C penalty term was accomplished by performing cross-validation using the scikit-learn stratified k-fold library. The default five splits were chosen, and the shuffle variable was set to true. The F1 macro average was selected to represent a model’s performance.

#### SGDP

The SVM library from scikit-learn was used to train the model for the SGDP dataset. Stratified k-fold cross-validation was performed using the standard scikit-learn library. Seven continental groups were predicted from this cohort (Africa, West Eurasia, East Asia, South Asia, Oceania, Central Asia Siberia, and America), as the subcontinental groups needed more samples per group to train an accurate model. The F1 macro average was chosen as a representation of a model’s performance to account for the imbalanced data.

## Results

### Model Performance

We report the performance of the gnomAD, 1kGP, and SGDP continental models using external validation sets (**Figure 2A-F**), and cross-validation results on the subcontinental models (**Figures S2 and S3**) were performed because additional datasets with the same subcontinental labels were not available.

**Figure 2.**
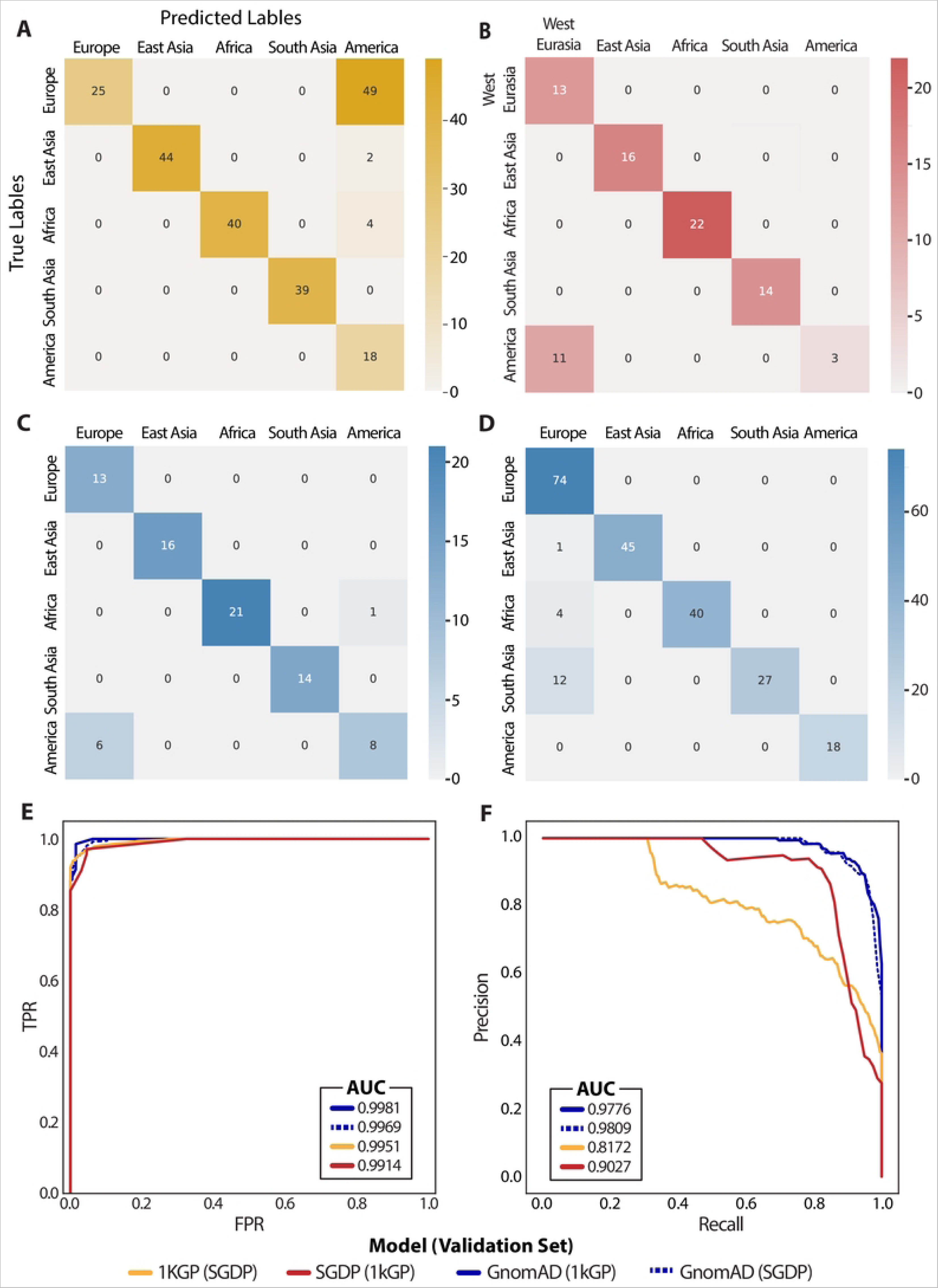
Continental ancestry inference model performance. A-D. Confusion matrices of the 1kGP model using SGDP as validation (A), SGDP model using 1kGP as validation (B), gnomAD model using 1kGP as validation (C), and gnomAD model using SGDP as validation (D). E. Macro-averaged ROC curves. F. Macro-averaged precision-recall curves.

Confusion matrices are shown in **Figures 2A-D**, providing the ancestry prediction for each sample in the validation data. In the 1kGP and SGDP models, we see some discrepancies between the European and American groups. In the case of the 1kGP model (**Figure 2A**), some SGDP samples labeled as European are predicted to be American. Similarly, in the SGDP model, some 1kGP samples labeled as American are predicted as European. This may be due to a higher similarity of the feature space between European and American samples than other groups (**Figure S3**). The gnomAD model is validated with 1kGP (**Figure 2C**) and SGDP (**Figure 2D**) samples. Overall, all continental models have a high area under the curve in both ROC (**Figure 2E**) and Precision-Recall (**Figure 2F**) curves, described in the figure legend.

The gnomAD East Asian and European subcontinental models have accuracies of 99.90% and 80.92%, respectively (**Figure S2A, B**). The results for the 1kGP subcontinental model are obtained by averaging the probabilities for each sample across cross-validation folds and then computing the confusion matrix (**Figure S4**). The accuracies for the 1kGP subcontinental models are as follows: Africa, 90.26%; America, 93.06%; East Asia, 87.23%; Europe, 94.29%; South Asia, 85.86%.

### Feature Interpretation

Feature importance for the gnomAD continental model was calculated using SHAP^37^ values to provide insight into which SNVs and their corresponding genes have the most impact on the model predictions. SHAP values for the 1kGP and SGDP models were not calculated because the memory requirement for the kernel explainer was too high due to the number of features in the models.

Global feature importance for the gnomAD continental model is reported by aggregating SHAP values across each gene and taking the mean absolute value of each gene across 2,800 of the training samples (**Figure S5**). The ‘knownCanonical’ genes table was downloaded from the UCSC Table Browser using assembly GRCh37 to get the genomic interval for each gene. If a region contains multiple genes, we combine the genes to form a non-overlapping genomic interval (e.g., ANKRD45, TEX50). Of the 77,402 variants used to train the model, 3,231 were not located in gene regions and were removed from further analysis. The most significant gene impacting the model is Keratin Associated Protein 19-8. Samples with a variant in this gene are more likely to be predicted as American.

We also aggregated SHAP values across larger cytolocations to visualize which regions across the genome are most impactful in the model predictions (accessed using this file: (https://hgdownload.soe.ucsc.edu/goldenPath/hg19/database/cytoBand.txt.gz). **Figure S6** shows the feature importance for an individual from the training data labeled as African. Regions are colored by population label with the maximum absolute SHAP value. Regions that have the most impact on predicting the sample African are ‘chromosome 1: 172,900,000-176,000,000’ and ‘chromosome 5: 63,200,000_66,700,000’.

### Comparison of Genetic vs. Self-Reported Ancestry in Clinical Samples

SNVstory was implemented on an in-house dataset of clinical exome sequencing testing from 293 individuals generated by the Institute for Genomic Medicine Clinical Laboratory to demonstrate the application of our models. We compare the model predictions to the self-reported ancestry of the proband (**Table S2**). Self-reported race is derived from the paternal/maternal ethnic background. Ethnicity is categorized into one of three groups: Non-Hispanic or Latino, Hispanic or Latino, and Unknown/Not Reported Ethnicity. Race is classified into one of five groups: White, Asian, Bi-racial/Multi-racial, Black or African American, and Unknown/Unspecified. Due to the broadness of these categories, we report the comparison between predicted genetic ancestry for the continental models only (**Table 1**).

**Table 1.**
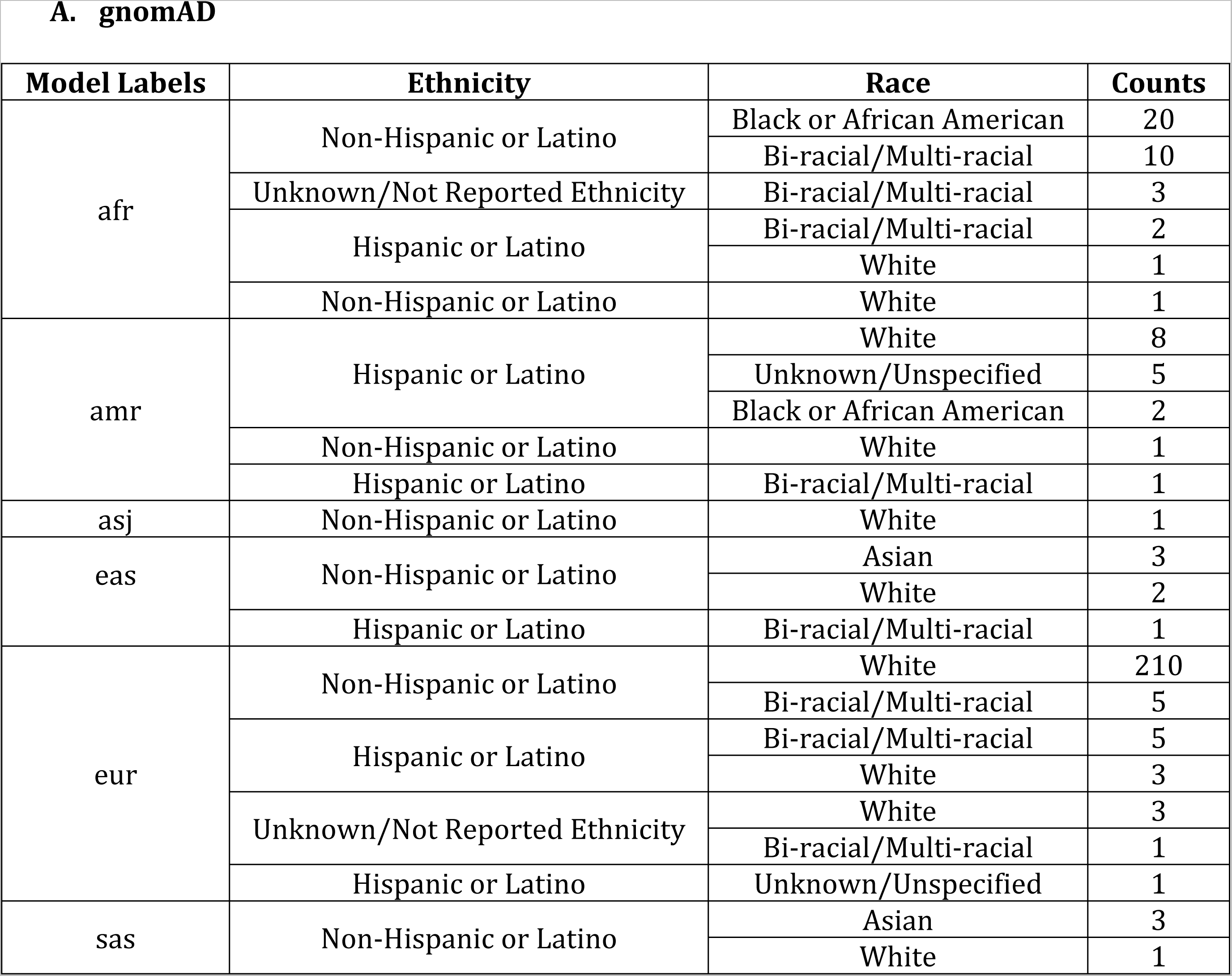
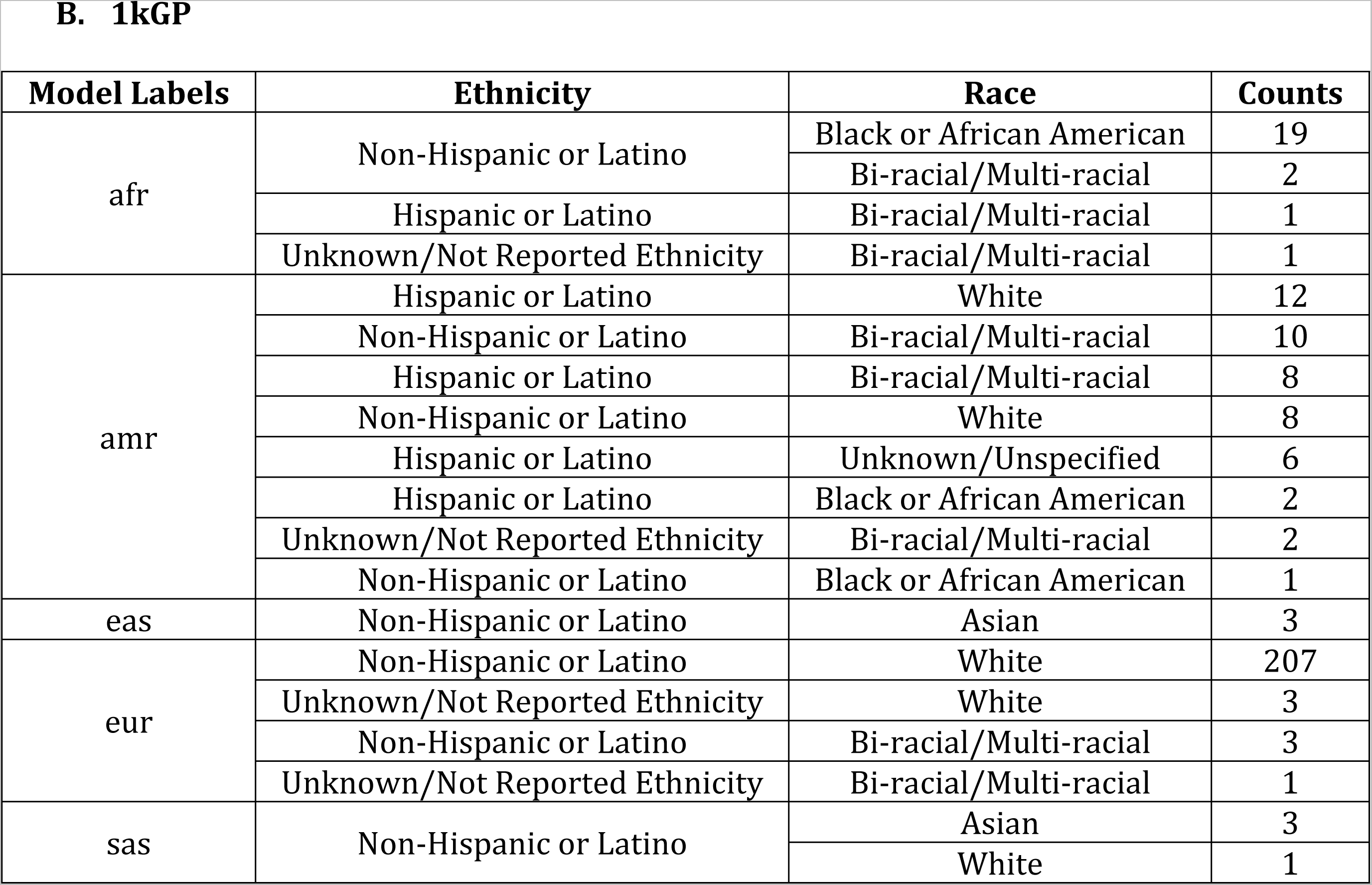
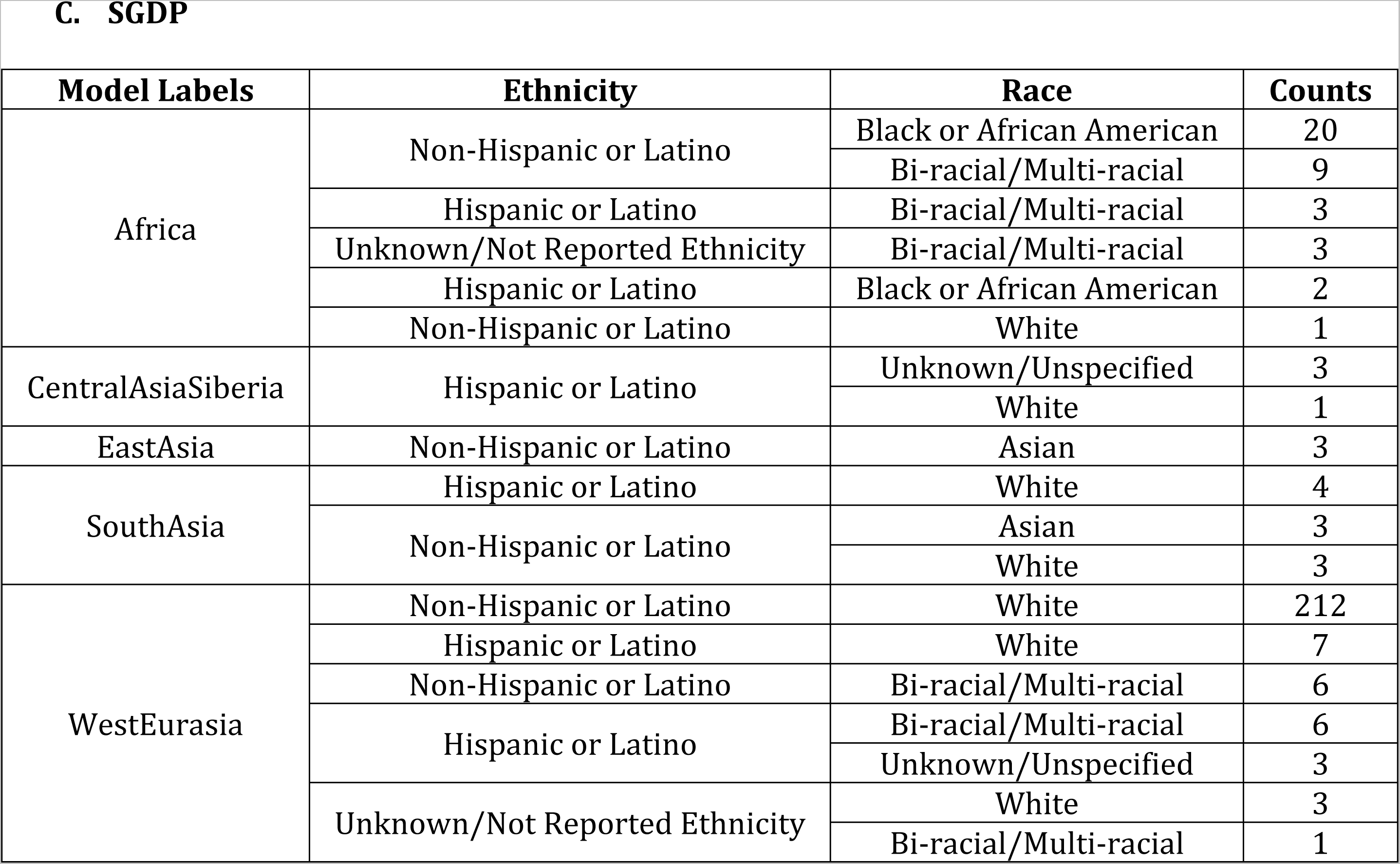
Genetic ancestry versus self-reported ethnicity and race. Value counts of genetic ancestry model predictions trained using gnomad **(A)**, 1kGP **(B)**, and SGDP **(C)** compared to self-reported ethnicity and race.

A. gnomAD
| Model Labels | Ethnicity | Race | Counts |
| --- | --- | --- | --- |
| afr | Non-Hispanic or Latino | Black or African American | 20 |
|  |  | Bi-racial/Multi-racial | 10 |
|  | Unknown/Not Reported Ethnicity | Bi-racial/Multi-racial | 3 |
|  | Hispanic or Latino | Bi-racial/Multi-racial | 2 |
|  |  | White | 1 |
|  | Non-Hispanic or Latino | White | 1 |
| amr | Hispanic or Latino | White | 8 |
|  |  | Unknown/Unspecified | 5 |
|  |  | Black or African American | 2 |
|  | Non-Hispanic or Latino | White | 1 |
|  | Hispanic or Latino | Bi-racial/Multi-racial | 1 |
| asj | Non-Hispanic or Latino | White | 1 |
| eas | Non-Hispanic or Latino | Asian | 3 |
|  |  | White | 2 |
|  | Hispanic or Latino | Bi-racial/Multi-racial | 1 |
| eur | Non-Hispanic or Latino | White | 210 |
|  |  | Bi-racial/Multi-racial | 5 |
|  | Hispanic or Latino | Bi-racial/Multi-racial | 5 |
|  |  | White | 3 |
|  | Unknown/Not Reported Ethnicity | White | 3 |
|  |  | Bi-racial/Multi-racial | 1 |
|  | Hispanic or Latino | Unknown/Unspecified | 1 |
| sas | Non-Hispanic or Latino | Asian | 3 |
|  |  | White | 1 |

**B. 1kGP**
| **Model Labels** | **Ethnicity** | **Race** | **Counts** |
| --- | --- | --- | --- |
| afr | Non-Hispanic or Latino | Black or African American | 19 |
|  |  | Bi-racial/Multi-racial | 2 |
|  | Hispanic or Latino | Bi-racial/Multi-racial | 1 |
|  | Unknown/Not Reported Ethnicity | Bi-racial/Multi-racial | 1 |
| amr | Hispanic or Latino | White | 12 |
|  | Non-Hispanic or Latino | Bi-racial/Multi-racial | 10 |
|  | Hispanic or Latino | Bi-racial/Multi-racial | 8 |
|  | Non-Hispanic or Latino | White | 8 |
|  | Hispanic or Latino | Unknown/Unspecified | 6 |
|  | Hispanic or Latino | Black or African American | 2 |
|  | Unknown/Not Reported Ethnicity | Bi-racial/Multi-racial | 2 |
|  | Non-Hispanic or Latino | Black or African American | 1 |
| eas | Non-Hispanic or Latino | Asian | 3 |
| eur | Non-Hispanic or Latino | White | 207 |
|  | Unknown/Not Reported Ethnicity | White | 3 |
|  | Non-Hispanic or Latino | Bi-racial/Multi-racial | 3 |
|  | Unknown/Not Reported Ethnicity | Bi-racial/Multi-racial | 1 |
| sas | Non-Hispanic or Latino | Asian | 3 |
|  |  | White | 1 |
| Africa | Non-Hispanic or Latino | Black or African American | 20 |
|  |  | Bi-racial/Multi-racial | 9 |
|  | Hispanic or Latino | Bi-racial/Multi-racial | 3 |
|  | Unknown/Not Reported Ethnicity | Bi-racial/Multi-racial | 3 |
|  | Hispanic or Latino | Black or African American | 2 |
|  | Non-Hispanic or Latino | White | 1 |
| CentralAsiaSiberia | Hispanic or Latino | Unknown/Unspecified | 3 |
|  |  | White | 1 |
| EastAsia | Non-Hispanic or Latino | Asian | 3 |
| SouthAsia | Hispanic or Latino | White | 4 |
|  | Non-Hispanic or Latino | Asian | 3 |
|  |  | White | 3 |
| WestEurasia | Non-Hispanic or Latino | White | 212 |
|  | Hispanic or Latino | White | 7 |
|  | Non-Hispanic or Latino | Bi-racial/Multi-racial | 6 |
|  | Hispanic or Latino | Bi-racial/Multi-racial | 6 |
|  |  | Unknown/Unspecified | 3 |
|  | Unknown/Not Reported Ethnicity | White | 3 |
|  |  | Bi-racial/Multi-racial | 1 |

Most of the individuals share agreement between genetic ancestry and ethnicity/race, e.g., for those predicted to be European, a match of White / Non-Hispanic or Latino for race /ethnicity occurs in 92.5%, 96.7%, and 89.1% of individuals by the gnomAD (**Table 1A**), 1kGP (**Table 1B**), and SGDP (**Table 1C**) models, respectively. However, several cases exist where individuals are self-reported as White while having a different genetic ancestry across multiple models, and vice versa. Additionally, 13 of our cases have either Unknown/Not Reported Ethnicity or Unknown/Unspecified Race. As discussed in the Introduction, the ability to refine or add genetic ancestry information in these cases is helpful for added diagnostic precision in variant filtering/prioritization.

### Model Interpretation for Indeterminant Samples

Most of our in-house dataset has agreement across all three continental models (81.9% of samples) and even more across at least two continental models (98.0%). A disproportionate number of individuals share disagreement across all three models between those that are self-reported as Bi-racial or Multi-racial vs. those that are White, Asian, Black or African American (50% vs. 9% disagreement, respectively). Those individuals with Unknown/Unspecified Race are not included in this calculation. These results suggest our models have worse performance on admixed samples, where two or more populations may be present. In reporting results, we use the label with the highest probability. Some discrepancies between model results may be mitigated by adding a minimum threshold on the probability required to obtain a result.

### Individualized Ancestry Report

Here, we illustrate the ability of SNVstory to provide ancestry predictions in an easily visualized format for individual samples (**Figure 3**). The probabilities for the gnomAD and the 1kGP continental models were 100% European, while the SGDP continental model was 95% West Eurasia. The gnomAD subcontinental model has the highest probability (48%) for North-Western European (nfe_nwe), and the 1kGP subcontinental model has the highest probability (100%) for British From England and Scotland (eur_gbr). The subcontinental model probabilities are weighted by the continental probabilities, which are returned as 0% probability for the remaining models. These predictions agree with the true sample ancestry taken from the 1kGP validation set.

**Figure 3.**
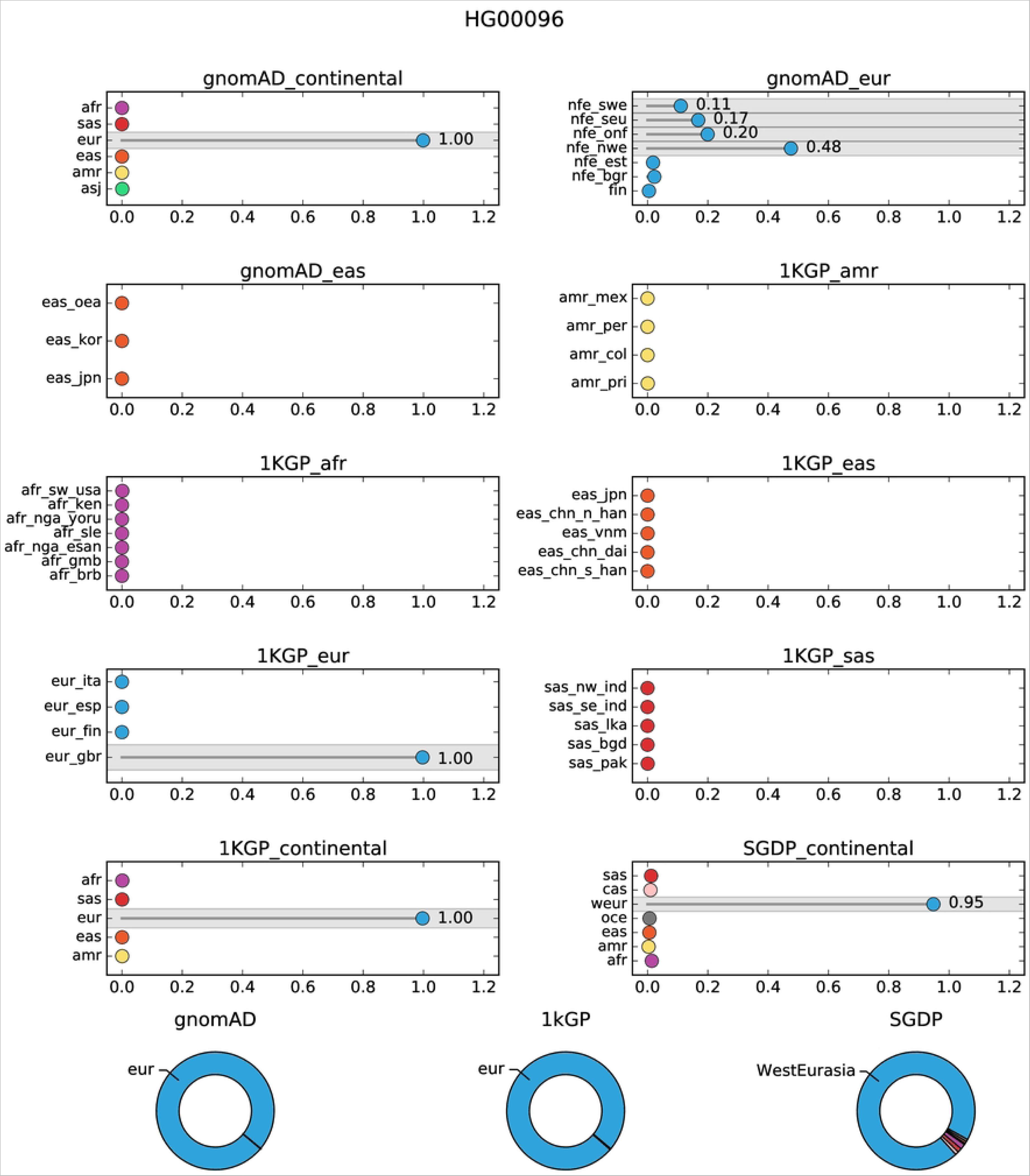
SNVstory ancestry report. The representative output of model results from SNVstory for a European sample taken from the 1kGP dataset.

## Discussion

We have described a method to predict ancestry from genomic data that provides multiple improvements over existing ancestry inference tools. Firstly, SNVstory incorporates samples/variants from three different curated datasets, expanding the number of labels and the granularity of the model classification beyond the main continental divisions. Secondly, drawing upon the gnomAD database produces a much larger number of variants on which our models were trained, providing the opportunity to classify ancestry on a wider (or more diverse) range of features. Thirdly, SNVstory excludes consanguineous samples from training, ensuring that the overrepresentation of closely related individuals does not bias the model. Finally, our novel implementation is optimized for individualized results rather than clustering large cohorts of samples into shared ancestral groups.

In our gnomAD model, we introduce a method to simulate individual samples from aggregate allele frequencies of a known population. This is potentially useful for any study requiring access to reference variants from a population where data from individual samples is obfuscated. One limitation in our approach is that we did not account for linkage disequilibrium between variants when simulating individual samples. This could result in some samples with patterns of variants that do not exist in actual samples. An improvement in future models would be to remove variants with high levels of linkage disequilibrium between them. If high recognizability to actual samples is required, established metrics of linkage disequilibrium, such as the correlation coefficient *r*^2^, could be used to measure the ‘realness’ of a simulated sample based on existing variant patterns, and simulated VCFs could be validated based on this quality. However, in practice, the larger pool of variants provided by gnomAD more than compensates for the lost dependence among proximal groups of variants. We have demonstrated that the performance of the gnomAD models with simulated individuals is comparable to that of models trained with actual samples.

With the growing number of reference datasets containing individuals from diverse ancestral backgrounds, it is possible to build ancestry prediction models that reflect these populations. However, there is room for improvement, as our most diverse dataset (SGDP) includes the fewest samples. We could not build subcontinental models as granular as the labels provided because there were as few as two samples per label for many instances. Additionally, our model cannot accurately predict ancestry proportions in samples with admixed ancestry. Most admixture prediction software depends on a priori knowledge of the number of non-admixed populations and requires representation from such populations. There is limited availability of reference samples from admixed individuals, so our training data lacked representation from any admixed samples. Efforts to expand the number of reference sequences for diverse and admixed populations will provide opportunities to fill this gap.

SNVstory’s feature-importance capacity is unique among ancestral tools and could have significant clinical utility. The clinical application of most ancestral prediction tools is limited to simply predicting the patient’s ancestry. However, SNVstory’s unique capability to describe a given locus as characteristic, or atypical, of a given ancestry could lead to improved prioritization of variants. For example, SNVstory finds the most ancestrally informative gene on average to be KRTAP19-8, which is greatly enriched for SNVs predictive of Native American/Latino ancestry (**Figure S5**). This gene is a known driver of thyroid lymphoma^38^, a disorder that is the second-most-common type of cancer among Hispanic women^39^ but not even among the top five cancer types among women worldwide^40^. The inferred distinctiveness of Latino copies of KRTAP19-8 suggests that rare founder mutations in this gene may contribute to increased rates of thyroid cancer among women of Hispanic ancestry. The ability to target variants in genes inherited from specific populations adds a new tool to the diagnostician’s toolkit and could lead to improved patient outcomes.

Finally, our approach allows users to reliably execute our models given a single-sample or multi-sample VCF, with results tailored toward ancestry assignment for an individual sample. This provides immediately useful ancestry information in the clinical setting, where ancestry can be used to inform diagnostic or therapeutic decisions. Specifically, a subject’s ancestry can be used to help prioritize variants that may be rare in one population but not another. In the clinical setting, it may be essential to recognize the difference between ethnicity, race, and genetic ancestry in determining the optimal therapy or drug dosage.

Given the widespread availability of genome sequencing data and models like SNVstory that can accurately predict ancestry, we advocate for genetic ancestry to become the standard classification reported for genetic studies and clinical applications, where appropriate. Genetic ancestry offers enormous advantages over other self-reported information, such as ethnicity or race, because it supplies biological characteristics of a population and is consistently measurable. This advantage will only increase as more populations are sequenced and ancestry prediction becomes more reliable, and we improve our ability to contextualize the impact of genetic ancestry on clinical decision-making.

## Acknowledgments

We thank the Nationwide Foundation Pediatric Innovation Fund for generously supporting this project.

## Author Contributions

AB and AR processed data and trained models for gnomAD, 1kGP, and SGDP. AR designed the methods to simulate data from gnomAD allele frequencies and cross-validation architecture. AB and DC prepared figures and tables. AB wrote the first draft of the paper. JG, DC, AR, and PW assisted in preparing or revising the paper. AR and AB wrote the SNVstory software package. PW and EM supervised the project.

## Data Availability

The training data for our model are available as follows. gnomAD v2.1 data is available from https://gnomad.broadinstitute.org/downloads/. 1000 Genomes Project data is shared via the International Genome Sample Resource and can be accessed from https://www.internationalgenome.org/data-portal/data-collection/30x-grch38. Simons Genome Diversity Project data is available from the European Nucleotide Archive under project PRJEB9586. SNVstory is an open-source model and is available from https://github.com/nch-igm/snvstory.

## Ethics Approval and Consent to Participate

This study was reviewed and approved by the Institutional Review Board (IRB) of The Abigail Wexner Research Institute at Nationwide Children’s Hospital (Office for Human Research Protections (OHRP) IORG0000326; IRB00000568) as IRB17-00206 (“Institute for Genomic Medicine Comprehensive Profiling for Cancer, Blood, and Somatic Disorders”). The participant’s legal guardian/next of kin provided written informed consent to participate in this study.

